# Crystal structure of chloroplast fructose-1,6-bisphosphate aldolase from the green alga *Chlamydomonas reinhardtii*

**DOI:** 10.1101/2021.12.28.474321

**Authors:** Théo Le Moigne, Edoardo Sarti, Antonin Nourisson, Alessandra Carbone, Stéphane D. Lemaire, Julien Henri

## Abstract

The Calvin-Benson cycle fixes carbon dioxide into organic triosephosphates through the collective action of eleven conserved enzymes. Regeneration of ribulose-1,5-bisphosphate, the substrate of Rubisco-mediated carboxylation, requires two lyase reactions catalyzed by fructose-1,6-bisphosphate aldolase (FBA). While cytoplasmic FBA has been extensively studied in non-photosynthetic organisms, functional and structural details are limited for chloroplast FBA encoded by oxygenic phototrophs. Here we determined the crystal structure of plastidial FBA from the unicellular green alga *Chlamydomonas reinhardtii* (Cr). We confirm that CrFBA folds as a TIM barrel, describe its catalytic pocket and homo-tetrameric state. Multiple sequence profiling classified the photosynthetic paralogs of FBA in a distinct group from non-photosynthetic paralogs. We mapped the sites of thiol- and phospho-based post-translational modifications known from photosynthetic organisms and predict their effects on enzyme catalysis.

## Introduction

Photosynthesis is a paramount biological chain reaction sustaining life on Earth through the conversion of light energy into chemical energy and the subsequent fixation of atmospheric carbon into triosephosphates (Pfannschmidt and Yang 2012). The Calvin-Benson cycle (CBC) is an eleven enzymes cycle resulting in the production of six molecules of glyceraldehyde-3-phosphate (G3P) from three ribulose-1,5-bisphosphate (RuBP) and three carbon dioxides (Bassham, Benson et al. 1950). In order to regenerate RuBP and sustain the cycle, five G3P are recycled into three RuBP by a series of reactions involving eight out of eleven CBC enzymes. Two G3P are converted to dihydroxyacetone phosphate (DHAP), one is reacted with G3P to form fructose-1,6-bisphosphate (FBP) and the other is ligated with erythrose-4-phosphate (E4P) to form sedoheptulose-1,7-bisphosphate (SBP) (Sharkey 2019). Both reactions are catalyzed by fructose-bisphosphate aldolase (FBA, E.C. 4.1.2.13), which therefore acts in two distinct steps of the CBC. Indeed, it can use two different couples of substrates and provides substrates to downstream fructose-1,6-bisphosphatase (FBPase) and sedoheptulose-1,7-bisphosphatase (SBPase). *Pisum sativum* FBAs were isolated from plant, and their enzymatic activity assayed (Anderson and Pacold 1972) (Razdan, Heinrikson et al. 1992). Change in FBA abundance are correlated with the activity of the CBC, supporting a potential control of FBA over photosynthetic carbon fixation (Haake, Zrenner et al. 1998) (Simkin, McAusland et al. 2015).

FBA is ubiquitous in the three phylogenetic kingdoms because it acts in the fourth step of glycolysis preparatory phase where it splits FBP into DHAP and G3P (Anderson and Advani 1970) (Fothergill-Gilmore and Michels 1993). The nuclear genome of the green alga *Chlamydomonas reinhardtii* codes for at least four aldolases paralogs: FBA1 (Uniprot entry A8JCY4, Ensembl gene <u>CHLRE_01g006950v5</u>), FBA2 (Uniprot entry A8I9H5, Ensembl gene CHLRE_02g093450v5), ALDCHL/FBA3 (Uniprot entry Q42690, Ensembl gene <u>CHLRE_05g234550v5</u>), and FBA4 (Uniprot entry A8I2Q9, Ensembl gene <u>CHLRE_02g115650v5</u>). ALDCHL/FBA3 is predicted to be localized in the chloroplast by its amino-terminal 17-residue (Tardif, Atteia et al. 2012) or 27-residue transit peptide (Almagro Armenteros, Salvatore et al. 2019). Mature proteins containing 333/323 residues was quantified as the 9^th^ most abundant protein in Chlamydomonas proteome (Schroda, Hemme et al. 2015), the third most abundant CBC enzyme after Rubisco large and small subunits (Hammel, Sommer et al. 2020). The estimated concentration of FBA3 is 4.2-10.1 amol/cell (Wienkoop, Weiss et al. 2010) (Hammel, Sommer et al. 2020), or 75.1-658.5 μmol/L in the chloroplast stroma (Mettler, Muhlhaus et al. 2014) (Hammel, Sommer et al. 2020). DHAP and G3P substrates are respectively concentrated at 0.2 mmol/L and 0.01 mmol/L in Chlamydomonas illuminated by 41 μmol photons/m^2^/s, and their concentrations increase at 0.7 mmol/L and 0.03 mmol/L under 145 μmol photons/m2/s. Substrate over enzyme ratios are low (0.4 G3P:FBA, 9 DHAP:FBA at 145 μmol photons/m2/s and 75.1 μmol/L enzyme, the lowest estimate of FBA3 concentration), indicating that FBA functions far from substrates saturation. This undersaturation of the active site is further confirmed by the relatively high K_M_ values reported for a bacterial FBA, which has Michaelis-Menten constants (K_M_) for DHAP and G3P of 0.095 mmol/L and 1.17 mmol/L, respectively (Rozova, Khmelenina et al. 2010). Moderate turnover numbers (k_cat_) were observed ranging from 0.2 to 65 sec^−1^. Although there is a lack of knowledge about the kinetic properties of the chloroplast isoforms, we can hypothesize that the catalytic capacity of FBA in the plastidial context is likely to exert a kinetic control over CBC metabolic flow.

FBA structures are classified into two groups, differentiated by their catalytic mechanism (Fothergill-Gilmore and Michels 1993) (Reyes-Prieto and Bhattacharya 2007). Class I aldolases form a Schiff-base intermediate with the substrate DHAP while class II aldolases are metal-dependent enzymes using divalent cations (Flechner, Gross et al. 1999). The two classes have been described as evolutionary independent (Marsh and Lebherz 1992). Despite folding as a similar α/β TIM barrel (SCOPe lineage c.1.10), class I and II aldolases have dissimilar quaternary structures, class I aldolases being homotetrameric and class II aldolase homodimeric.

Mounting evidence supports that CBC enzymes are target of multiple post-translational modifications by means of reversible oxidation of cysteine thiols (reviewed in (Michelet, Zaffagnini et al. 2013) (Zaffagnini, Fermani et al. 2019) (Müller-Schüssele, Bohle et al. 2021)) or threonine/serine phosphorylation ((Wang, Gau et al. 2014) (Werth, McConnell et al. 2019). CrFBA3 has been retrieved in the thioredoxome of *Chlamydomonas reinhardtii* supporting the relative affinity of FBA3 for thioredoxin through residue Cys58 (Lemaire, Guillon et al. 2004) (Perez-Perez, Mauries et al. 2017). CrFBA3 was identified as a putative target of GSSG-mediated S-glutathionylation (Zaffagnini, Bedhomme et al. 2012), and it was also found undergoing S-nitrosylation on three cysteine sites (Cys58, Cys142, and Cys256 (Morisse, Zaffagnini et al. 2014), and phosphorylation on Ser54, Thr57, Ser64, Ser170, Ser176 (Wang, Gau et al. 2014) (Werth, McConnell et al. 2019). Moreover, FBA was reported to be modified by lysine crotonylation in *Carica papaya* (Liu, Yuan et al. 2018). FBA is subjected to lysine acetylation in *Arabidopsis* (Finkemeier, Laxa et al. 2011), while *Chlamydomonas* FBA was found to interact with CP12, a small intrinsically disordered protein specifically involved in the formation of inhibitory supercomplex with two CBC enzymes, namely phosphoribulokinase and glyceraldehyde-3-phosphate dehydrogenase (PRK and GAPDH, respectively) (Erales, Avilan et al. 2008) (Yu, Xie et al. 2020). Post-translational modifications and protein-protein interactions may support reversible regulatory mechanisms of protein catalysis but their interpretation requires a precise mapping of modified groups in the context of the protein structure.

A functional classification of the CrFBA paralogs among their homologs could support the unique role of CrFBA3 in the carbon fixation pathway. The sequence-based computational method ProfileView (Vicedomini, Bouly et al. 2019) has been recently designed to address the functional classification of the great diversity of homologous sequences hiding, in many cases, a variety of functional activities that cannot be anticipated. ProfileView relies on two main ideas: the use of multiple probabilistic models whose construction explores evolutionary information in large datasets of sequences (Bernardes, Zaverucha et al. 2016) (Ugarte, Vicedomini et al. 2018) (Vicedomini, Blachon et al. 2021) (Fortunato, Jaubert et al. 2016) (Amato, Dell’Aquila et al. 2017), and a new definition of a representation space where to look at sequences from the point of view of probabilistic models combined together. ProfileView has been previously applied to classify families of proteins for which functions should be discovered or characterized within known groups (Vicedomini, Blachon et al. 2021). It was proven very successful in identifying functional differences between otherwise phylogenetically similar sequences. Here, we meet a new challenge and use ProfileView to distinguish paralogs by their function in the FBA family.

In order to determine the structure-function-regulation relationships of photosynthetic FBA, we solved the high-resolution structure of the recombinant protein from Chlamydomonas, and performed a computational analysis that confirms functional segregation of CrFBA3 relative to the other FBA paralogs present in *Chlamydomonas*.

## Results

### CrFBA3 general structure

The recombinant FBA3 from C. reinhardtii was purified to homogeneity and the crystal structure was solved at 2.36 Å resolution. The protein crystallized in the C2 space group (table 1) and the crystallographic model includes eight polypeptide chains of 329-332 residues numbered according to the UniProtKB entry Q42690. Residual electron density at 1 σ additionally allowed us to place 29 sulfate ions, one chloride ion, and 989 water molecules in the crystal asymmetric unit. Amino acids prior to Met27 and after Thr360 have not been modelled because of a lack of electron density, as expected for peptidic extensions of low complexity. Alignment of the eight chains onto each other showed carbon alpha RMSD values comprised between 0.177 Å and 0.295 Å. All chains are therefore virtually identical except for some amino acid positions at N-terminal and/or C-terminal extremities, so we will focus on chain C for the following description, this chain being the most complete among the eight. Structural alignment of CrFBA3 to the protein data bank archive returned 470 matches, with highest similarities towards *Arabidopsis thaliana* glycolytic FBA (PDB ID: 6RS1, Q-score = 0.88; RMSD = 0.93 Å), with *Plasmodium falciparum* FBA (PDB ID: 4TR9, Q-score = 0.87; RMSD = 0.89 Å), and with *Toxoplasma gondii* FBA (PDB ID: 4D2J, Q-score = 0.87; RMSD= 0.95 Å). CrFBA3 is an α/β protein composed of eleven α-helices and nine β-strands folded as a TIM barrel (CATH classification: 3.20.20) and numerated from H1 to H11 and S1 to S9, respectively (figure 1). Regions of the protein with the largest chain-to-chain differences are otherwise highlighted by the highest B-factors, and are mainly localized on the H2, H3, H4, H5 and H10 and H11 α-helices and the loops that connect them (figure 2). These regions are located close to the active site of the protein and may allow rearrangements of its accessibility, in order to potentially accommodate substrates and products according to an induced fit mechanism.

**Table 1.** X-ray diffraction and crystallographic model statistics.

|  | **Fructose biphosphate aldolase** |
| --- | --- |
| **Wavelength** | 0.983995275547 |
| **Resolution range** | 46.67-2.36 (2.44-2.36) |
| **Space group** | C2 |
| **Unit cell** | 109.49-251.07-126.411<br>90-90.11-90 |
| **Total reflections** | 988770 (90253) |
| **Unique reflexions** | 138497 (13155) |
| **Multiplicity** | 7.1 (6.9) |
| **Completeness (%)** | 99.42 (94.42) |
| **Mean I/sigma(I)** | 13.34 (1.46) |
| **Wilson B-factor** | 40.19 |
| **R-merge** | 0.2 (1.495) |
| **R-meas** | 0.2159 (1.617) |
| **R-pim** | 0.08067 (0.6089) |
| **CC1/2** | 0.995 (0.566) |
| **CC\*** | 0.999 (0.85) |
| **Reflections used in refinement** | 138424 (13104) |
| **Reflections used for R-free** | 1984 (189) |
| **R-work** | 0.2025 (0.3002) |
| **R-free** | 0.2500 (0.3513) |
| **CC (work)** | 0.954 (0.775) |
| **CC (free)** | 0.921 (0.643) |
| **RMS (bonds)** | 0.004 |
| **RMS (angles)** | 0.63 |
| **Ramachandran favoured (%)** | 96.28 |
| **Ramachandran allowed (%)** | 3.34 |
| **Ramachandran outliers (%)** | 0.38 |
| **Rotamer outliers (%)** | 0.43 |
| **Clashscore** | 6.97 |
| **Average B-factor** | 46.74 |

**Figure 1.**
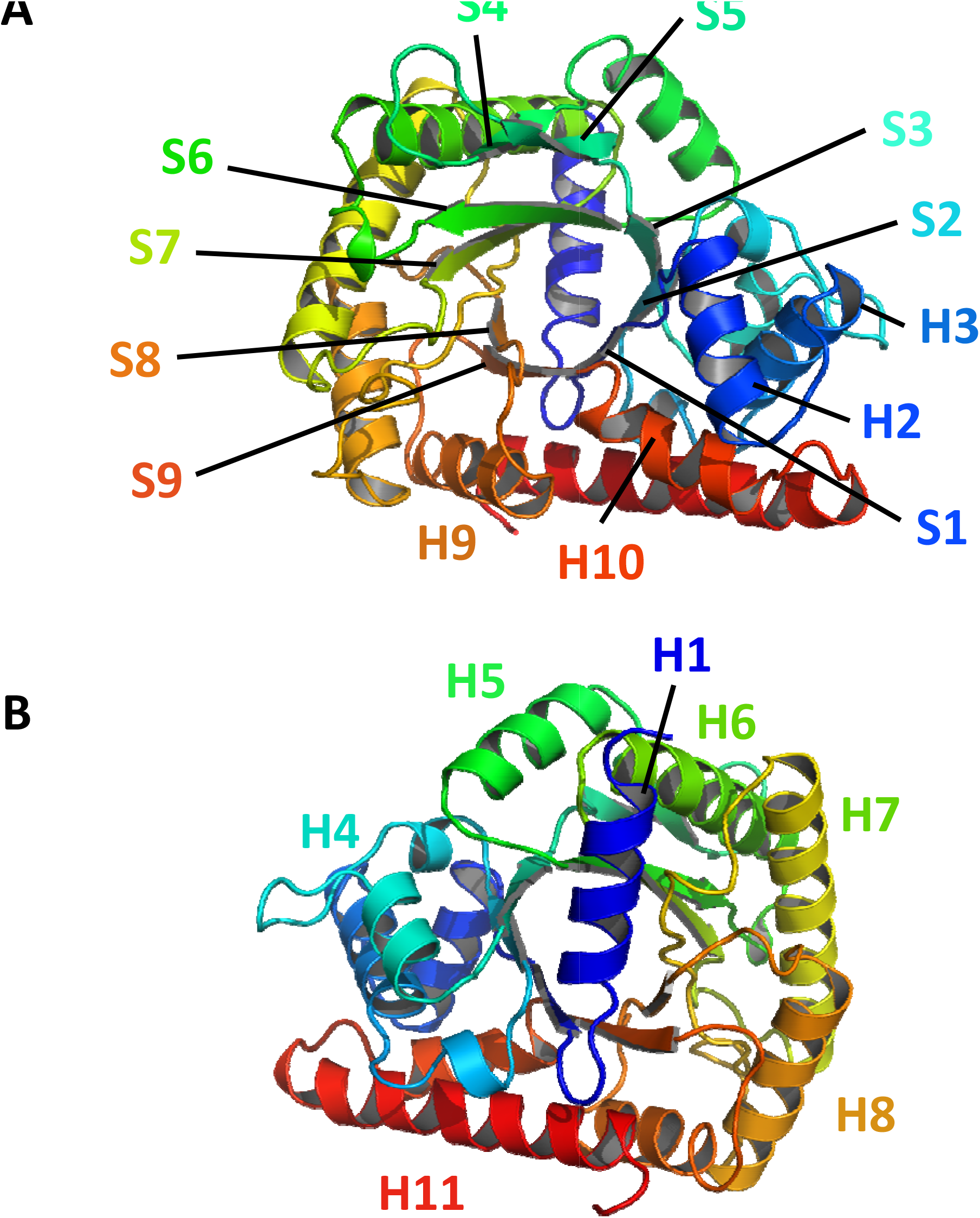
FBA topology. **A**. Front view of CrFBA1 represented as cartoon and coloured from N-terminal to C-terminal respectively from blue to red. Alpha helices and beta strand are indicated and named from H1 to H11 and from S1 to S9. **B**. Back view (180° rotation) of CrFBA1 with the same representation, coloration and indication as in A.

**Figure 2.**
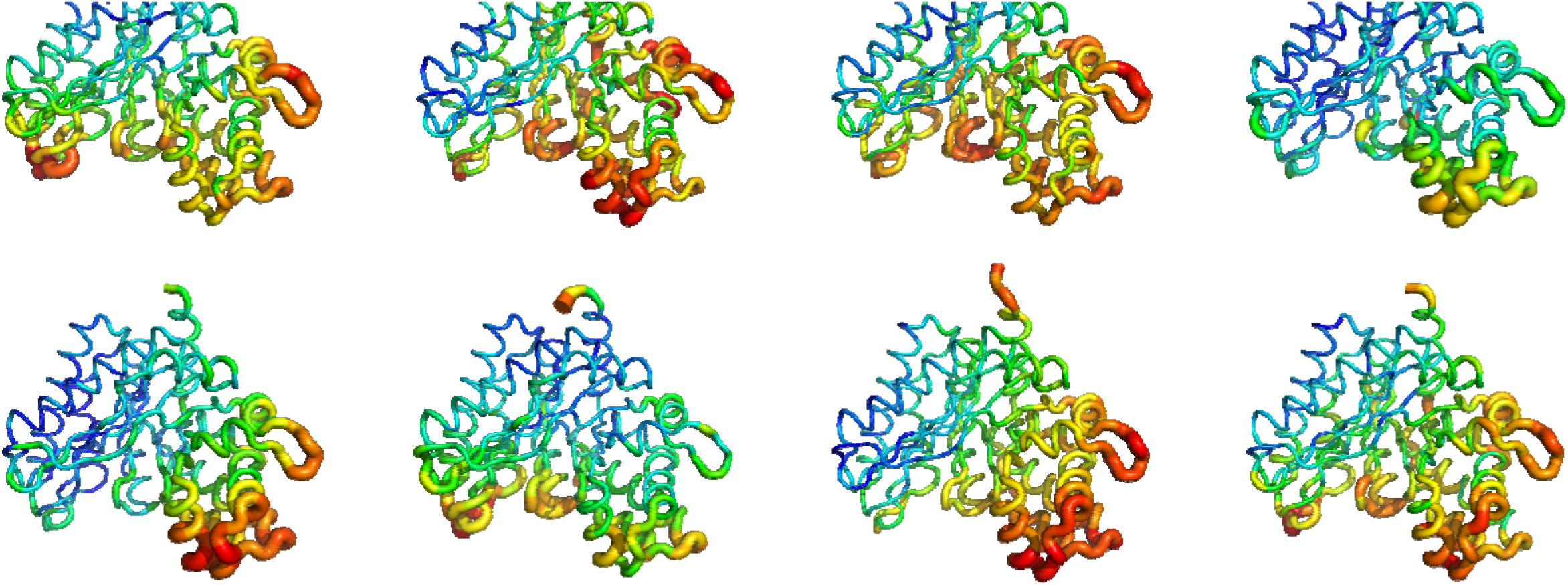
FBA local disorder. Representation of the eight monomeric chains of CrFBA1 of the asymmetric unit. Each amino acids is coloured from blue to red and is thicker in function of their B-factor value, respectively with the lower B-factor being blue and small and the higher being thick and red.

### CrFBA3 active site

Active site of CrFBA3 is localized in the center of the TIM barrel at the C-terminal pole of the β-strands. This site is a strongly electropositive pocket, containing one or two sulfate ions captured in the crystallization solution and several water molecules (figure 3). Active site of class I aldolase have already been described structurally and chemically in various organisms (Dalby, Dauter et al. 1999) (St-Jean, Lafrance-Vanasse et al. 2005) (Lafrance-Vanasse and Sygusch 2007) (Gardberg, Abendroth et al. 2011) (Gardberg, Sankaran et al. 2011). Eleven catalytic residues described in these articles are present in *Chlamydomonas reinhardtii* FBA3 at the following positions: Asp52, Ser54, Thr57, Lys125, Lys164, Arg166, Glu204, Glu206, Lys246, Ser288, and Arg318. Side chains of all catalytic residues are identically oriented placed in *Chlamydomonas reinhardtii* and mammals orthologs taken as references (supplementary figure 2). Human FBA has been described to perform its lyase activity by forming a Schiff base between the substrate and the lysine in position 229 while being stabilized by the two positively charged amino acids (Arg148 and Lys146) (Gamblin, Davies et al. 1991). The hypothesis of a similar catalytic mechanism between CrFBA3 and other FBA is comforted by superposing CrFBA3 structure to two FBA structures co-crystallized with their substrate in which amino acids from CrFBA3 are well superposed with those found in homologs from *Homo sapiens* FBA (HsFBA) (PDB ID: 4ALD) and *Oryctolagus cuniculus* FBA (OcFBA) (PDB ID: 1ZAI) with RMSD of 0.813 Å and 0.886 Å, respectively (figure 3).

**Figure 3.**
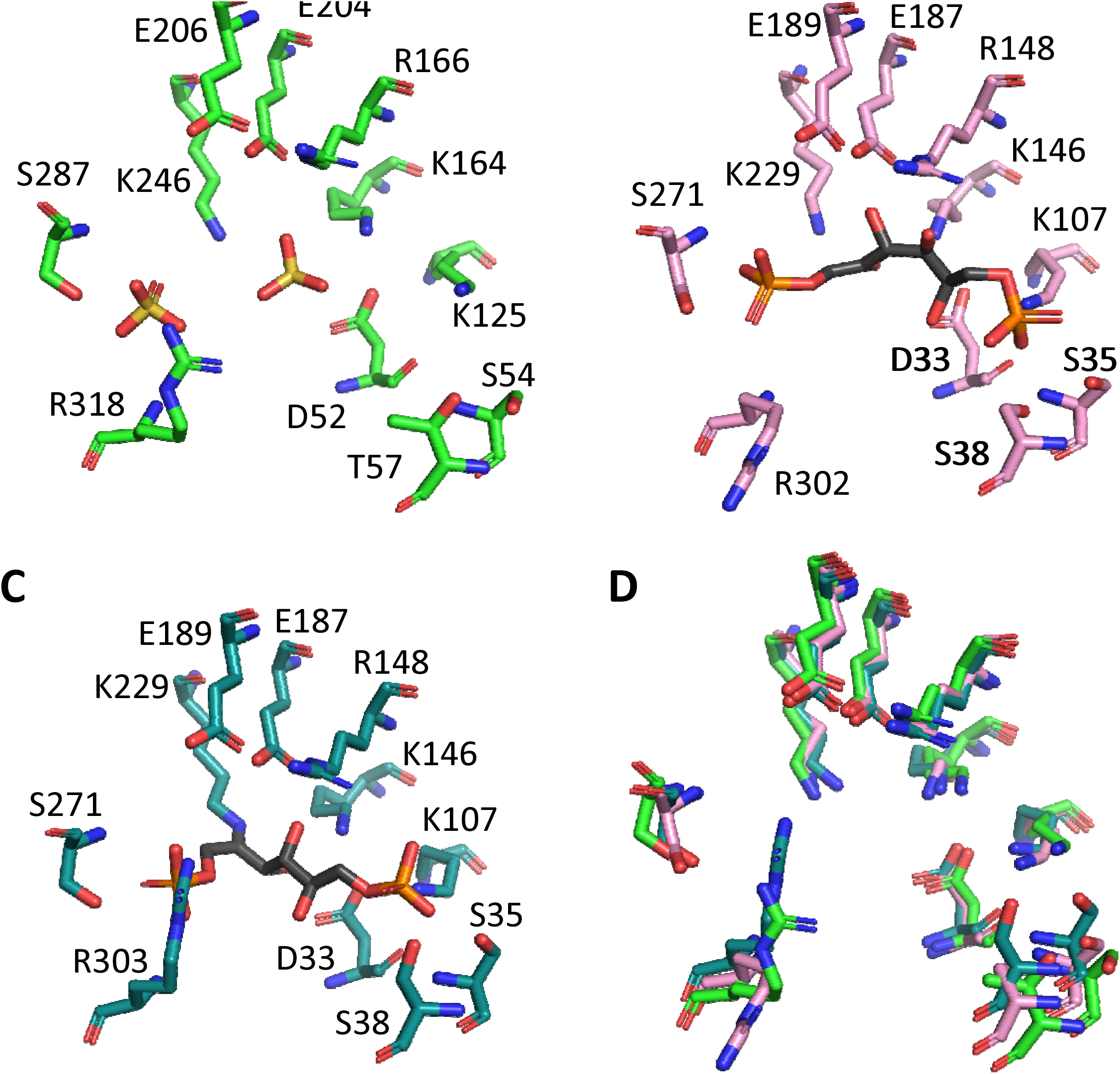
FBA active site. **A**. Stick representation of amino acids of the catalytic site of CrFBA1. Atoms are coloured with the CPK colouring code with an exception of carbon which is coloured in green. Two sulphate ions inside the pocket are also represented. **B**. Stick representation of amino acids of the catalytic site of HsFBA and a fructose-1,6-bisphosphate molecule from the PDB structure 4ALD aligned on the CrFBA1 structure. Atoms are coloured as in A. with carbon in light pink. **C**. Stick representation of amino acids of the catalytic site of OcFBA and a fructose-1,6-bisphosphate molecule from the PDB structure 1ZAI aligned on the CrFBA1 structure. Atoms are coloured as in A. with carbon in teal. **D**. Superposition of the amino acids from A. B. and C. coloured as in A. B. and C.

**Figure 4.**
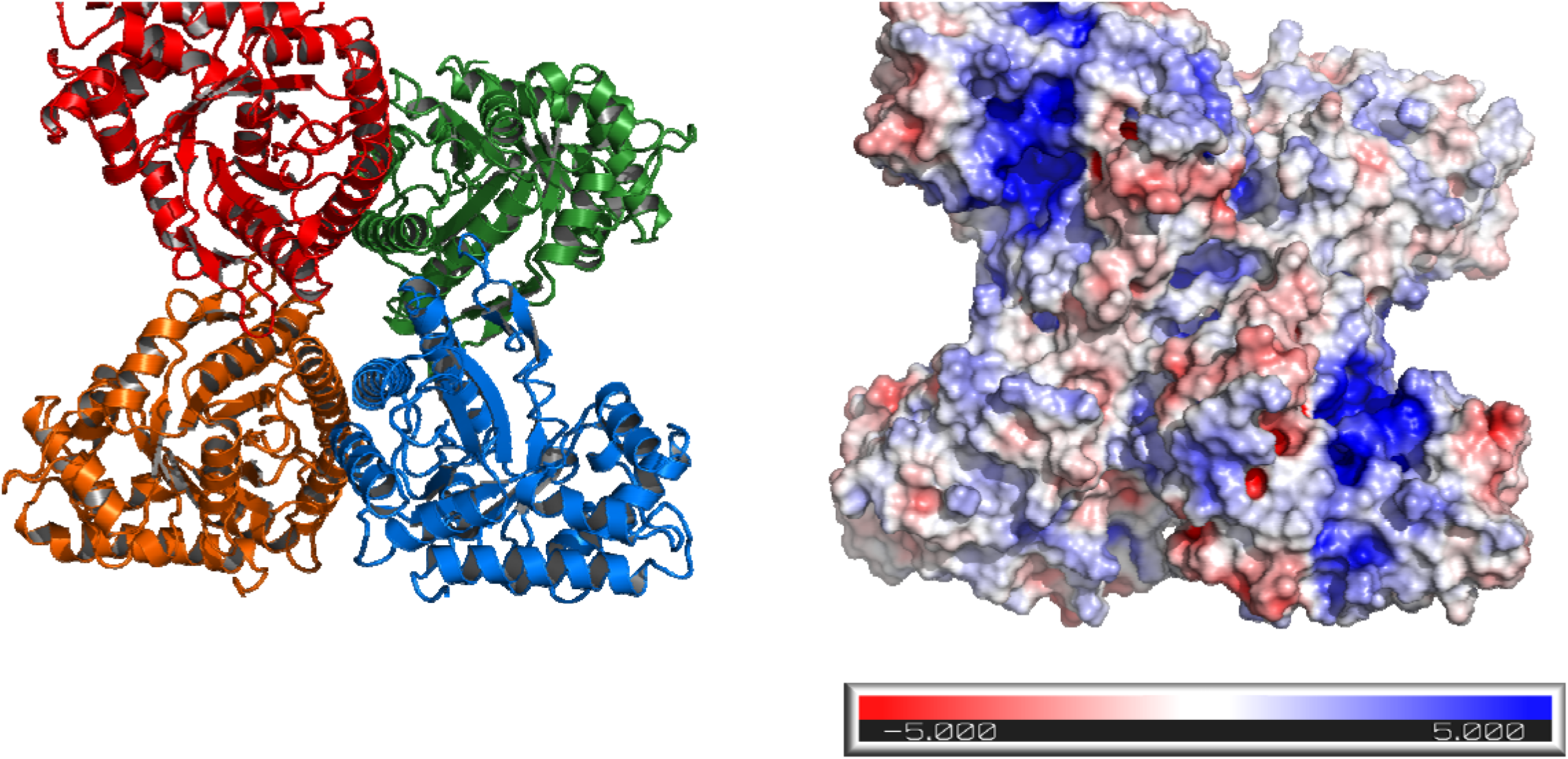
FBA quaternary structure. **A**. Cartoon representation of the homotetramer of CrFBA1. The different subunits are represented in red, green, orange and blue. **B**. Electrostatic surface of CrFBA1 calculated by PyMOL APBS and represented in a gradient from blue (electropositive) to red (electronegative). CrFBA1 is in the same orientation as in A.

### CrFBA3 is a homo-tetramer

CrFBA3 crystal has an asymmetric unit containing eight quasi-identical protein chain. The analysis of the quaternary structure of recombinant CrFBA3 protein indicated a homo-tetrameric state in solution, as the principal peak of protein elute at an approximative mass of 160 kDa closely matching the mass of four monomers of 39 kDa each (supplementary figure 1). Analytical size exclusion chromatography coupled to small angle X-ray scattering (SEC-SAXS) experiments confirmed the tetrameric state in solution (supplementary figure 2). Alteration of the redox state of CrFBA3 induced by oxidizing or reducing treatments prior to SEC-SAXS slightly impacted the estimated mass of the protein in solution being 142.15 kDa and 162.65 kDa for oxidized and reduced protein, respectively, and supporting a more compact state for oxidized FBA. Altogether, these results agree for a homo-tetrameric state of CrFBA3 in solution, which is also described in other structures of class I FBA in other species (Marsh and Lebherz 1992). In order to evaluate the physiological oligomeric state of FBA3, we fractionated soluble extracts of *Chlamydomonas* cultures over a calibrated size-exclusion column and the presence of FBA3 in the eluted fractions was monitored by western blot using anti-FBA3 antibodies. FBA3 eluted in one principal species with an apparent molecular weight of 170 kDa, consistent with that of a homotetramer.

### CrFBA3 sites of post-translational modifications

CrFBA3 possesses six cysteines in its mature sequence. At least three cysteines may be targeted for redox post-translational modifications mediated by thioredoxins (Perez-Perez, Mauries et al. 2017). The two couples of cysteines C142-C183 and C218-C256 are close enough, with respectively 8.3 Å and 10.2 Å between the two sulphur atoms, to form disulphide bonds. Recent redox-based proteomic studies identified Cys58, Cys142, and Cys256 as targets of S-nitrosylation while Cys58 and Cys142 were shown to be S-glutathionylated (Zaffagnini, Bedhomme et al. 2012). Despite being identified as putative redox target, Cys58 lateral chain is oriented toward the core of the protein and is not exposed to solvent (figure 5), raising the hypothesis that redox alteration of the thiol group of this residue demands local conformational changes or alternatively occurs in a later unfolded state of the protein. Also, Cys58 redox modification could be a co-translational modification rather than a post-translational modification occurring during the protein folding process. Cys142 is close to the catalytic cleft and to the external shell of the protein but is still poorly accessible to the solvent on our structure unless CrFBA3 undergoes to a dynamic remodelling of β-strands 4 and 5 (figure 1). These β-strands are actually the ones with the highest B-factors (figure 2), supporting the hypothesis of a rearrangement of this part of the protein in function of the redox conditions. Cys256 is actually the most exposed thiol of CrFBA3, making it the best candidate for a redox control of the enzyme. However, this residue is located at 19.6 Å from the catalytic Arg166 (fig. 5). Such a distance prevents a direct effect of redox modification on protein catalysis but could induce local structural shifts that could alter substrate binding and/or substrate processing.

**Figure 5.**
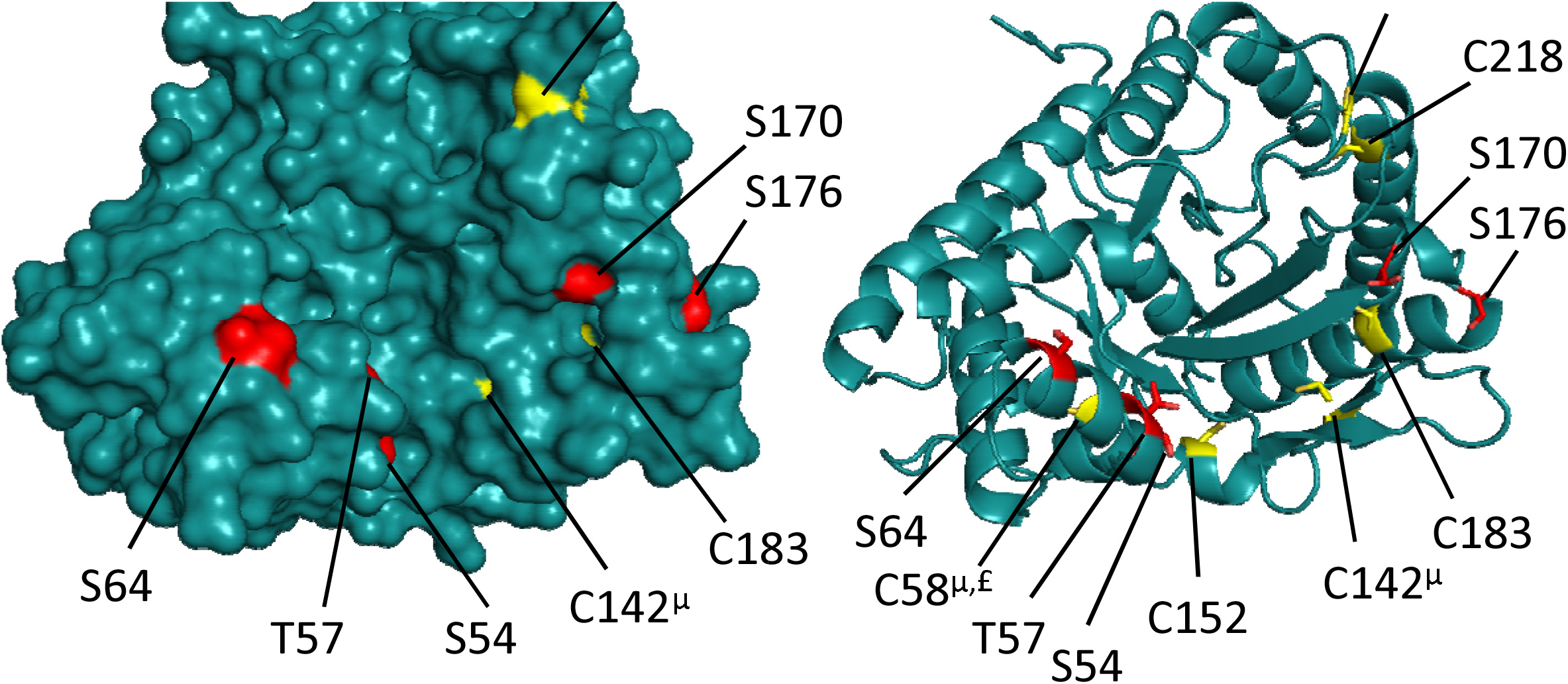
FBA redox modification + sites phosphorylation sites. A.B. nytrosylation=μ, £=gluthationylation,

Phosphoproteomic studies found five amino acids that are phosphorylated in CrFBA3 (Wang, Gau et al. 2014) (Werth, McConnell et al. 2019). These amino acids are dispersed in the primary structure, namely Ser54, Thr57, Ser64, Ser170, and Ser176. All corresponding sidechains are solvent exposed in CrFBA3 crystal structure (figure 5). Ser54, Thr57, and Ser64 are localized on the α-helix 2 which is a key element of the catalytic pocket. In fact, Ser54 and Thr57 are part of the amino acids needed to accommodate the substrate in the active site (Dalby, Dauter et al. 1999) (St-Jean, Lafrance-Vanasse et al. 2005) (Gardberg, Abendroth et al. 2011). Phosphorylation of these residues could interfere with the fixation of the substrate both via steric and charge obstructions, supporting a negative regulatory mechanism widely conserved as these residues are conserved from photosynthetic organisms to animals, archaea, and bacteria (supplementary figure 3). Residues Ser170 and Ser176 are localized on the loop between β-strand 6 and α-helix 6 separated by a 3^10^ helix. This loop is on the other side of the catalytic site, bordering it. While Ser176 is conserved in FBA from all living organisms, Ser170 is only conserved in photosynthetic organisms.

### CrFBA3 is recognized as the Calvin-Benson paralog in *Chlamydomonas reinhardtii*

Starting from a set of 2601 homologous sequences from both prokaryotic and eukaryotic photosynthetic organisms, we used ProfileView to construct a functional tree (figure 6) organizing the input sequences by function (see Methods), where, intuitively, distinguished subtrees of sequences correspond to distinguished functions. In order to functionally interpret ProfileView distinguished subtrees we used available functional knowledge on FBA homologous sequences included in the tree and belonging to other species (*Volvox carteri, Arabidopsis thaliana, Synechocystis sp. PCC 6803*, see Table 2) and checked whether Calvin-Benson paralogs were grouped under a common subtree or not. As shown in Figure 6A, ProfileView reconstructed three different subtrees, colored green, yellow and red. Calvin-Benson and non-Calvin-Benson sequences are located in different subtrees: the green one collects all known Calvin-Benson sequences, whereas the yellow and red ones collect all known non-Calvin-Benson ones. The two, yellow and red, subtrees contain almost exclusively sequences from unicellular and multicellular organisms, respectively. This means that the CrFBA3 sequence is neatly separated from its paralogs CrFBA1, CrFBA2 and CrFBA4: FBA3 appear in the Calvin-Benson subtree (figure 6B), whereas the others appear in the unicellular non-Calvin-Benson subtree (figure 6C).

**Figure 6.**
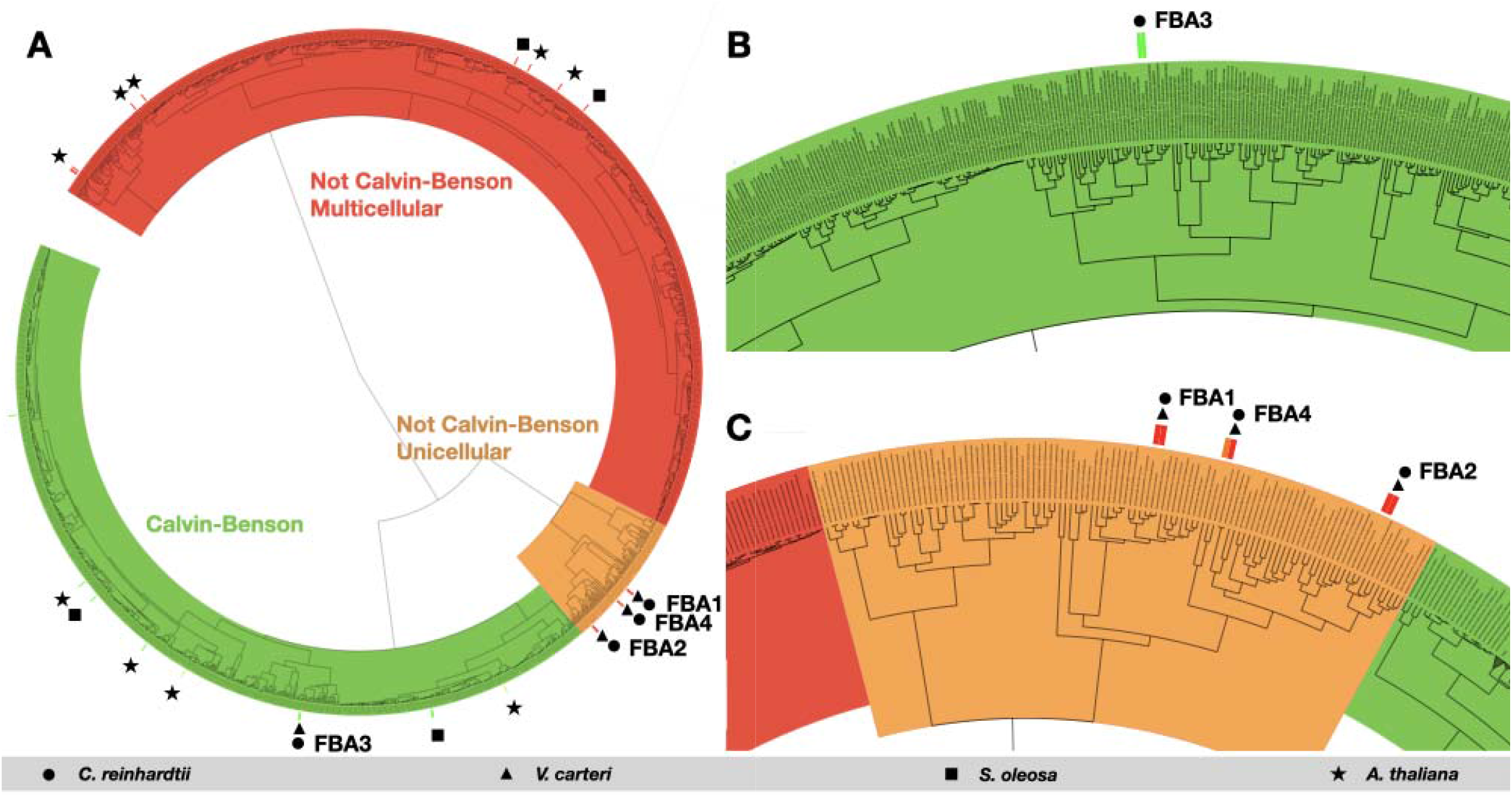
FBA functional tree. ProfileView tree showing functional relationship between FBA sequences of photosynthetic organisms: A) The tree separates FBA paralogs participating in Calvin-Benson (green subtree) and not participating in Calvin-Benson (orange and red subtrees, for unicellular and multicellular organisms respectively). For *C. reinhardtii*, the paralog FBA3 is localized in the Calvin-Benson subtree, whereas the other three paralogs FBA1, FBA2, FBA4 are grouped together in a different subtree. See Figure S4 for a high-resolution version to explore taxa. B) Zoom on the Calvin-Benson subtree containing the *C. reinhardtii* FBA3 sequence. C) Detail of the Unicellular Non-Calvin-Benson subtree, where the other *C. reinhardtii* FBA paralogs are found. In all panels, circles correspond to *C. reinhardtii* sequences, triangles to *V. carteri*, squares to *S. oleosa*, and stars to *A. thaliana*.

**Table 2.** Uniprot codes and the corresponding classification found by crossreferencing online databases. Classification was manually conducted by considering that Calvin-Benson FBA bear a chloroplast-addressing peptide, as predicted by ChloroP (Emanuelsson Nielsen et al. 1999) and Predalgo (Tardif et al. 2012), and present a circadian expression available for *Chlamydomonas* (Strenkert Merchant et al. PNAS 2019) and *Arabidopsis* (Romanowski et al. 2020).

| Uniprot code | Uniprot accession numbers | Classification |
| --- | --- | --- |
| Q42690_CHLRE | Q42690, A8JE10, Q36725 | *C. reinhardtii* FBA3<br>(positive test) |
| A8JCY4_CHLRE | A8JCY4 | *C. reinhardtii* FBA1<br>(negative test) |
| A8I9H5_CHLRE | A8I9H5 | *C. reinhardtii* FBA2<br>(negative test) |
| A8I2Q9_CHLRE | A8I2Q9 | *C. reinhardtii* FBA4<br>(negative test) |
| A0A0K9Q9C0_SPIOL | A0A0K9Q9C0 | Non-Calvin-Benson |
| A0A1P8B8S7_ARATH | A0A1P8B8S7 | Non-Calvin-Benson |
| ALFP2_ARATH | Q944G9, Q5XEU6, Q9SVJ6 | Non-circadian<br>chloroplastic |
| A0A1P8B8Q8_ARATH | A0A1P8B8Q8 | Non-Calvin-Benson |
| ALFC5_ARATH | O65581 | Non-Calvin-Benson |
| ALFC8_ARATH | Q9LF98 | Non-Calvin-Benson |
| A0A0K9QNV9_SPIOL | A0A0K9QNV9 | Non-Calvin-Benson |
| D8TKY4_VOLCA | D8TKY4 | Circadian chloroplastic |
| ALFC7_ARATH | P22197, O65582, Q53YH0 | Non-Calvin-Benson |
| D8UAC8_VOLCA | D8UAC8 | Non-Calvin-Benson |
| ALFP1_ARATH | Q9SJU4, Q2V473, Q93WF5, Q94C97 | Circadian chloroplastic |
| F4IGL5_ARATH | F4IGL5 | Circadian chloroplastic |
| F4IGL7_ARATH | F4IGL7 | Circadian chloroplastic |
| ALFP3_ARATH | Q9ZU52 | Non-circadian<br>chloroplastic |
| A0A0K9R8B2_SPIOL | A0A0K9R8B2 | Chloroplastic |
| D8U593_VOLCA | D8U593 | Non-Calvin-Benson |
| ALFC4_ARATH | F4KGQ0, Q8LET3, Q8W4G8, Q9LZR9 | Non-Calvin-Benson |
| D8UIJ0_VOLCA | D8UIJ0 | Non-Calvin-Benson |
| A0A0K9QFF9_SPIOL | A0A0K9QFF9 | Chloroplastic |
| F4JUJ5_ARATH | F4JUJ5 | Non-circadian<br>chloroplastic |
| A0A0K9QU20_SPIOL | A0A0K9QU20 | Chloroplastic |
| ALFC6_ARATH | Q9SJQ9 | Non-Calvin-Benson |
| B3H6D7_ARATH | B3H6D7 | Non-Calvin-Benson |
| A0A1P8BBB5_ARATH | A0A1P8BBB5 | Non-Calvin-Benson |
| A0A0K9R0B2_SPIOL | A0A0K9R0B2 | Chloroplastic |

## Conclusion

We describe here the structure of chloroplastic FBA3 from *Chlamydomonas reinhardtii*. If nothing seems different in the catalytic mechanism of CrFBA3 in comparison with already described class I FBA from other species, post-translational regulatory target sites of CrFBA3 appear original and chloroplast-specific. The two Cys58 and Cys142, both target of S-glutathionylation and S-nitrosylation, are only conserved in photosynthetic organisms and therefore could be specific targets of a distinct and stress-related regulation of CBC. The amino acid Ser170, target of phosphorylation events, is in the same case as it is conserved only in photosynthetic organism. Alpha helix 2 contain residues Ser54, Thr57, Cys58, and Ser64 that are subject to post-translational modifications, it also has high B-factor in comparison to the other secondary elements and is close to the active site. These characteristics making it a probable regulation cluster and an important area in the activity of CrFBA3 (Seoane and Carbone 2021).

This work also provides a computational analysis that allows a functional classification of the CrFBA paralogs among their homologs. This classification supports the unique role of CrFBA3 in the Calvin-Benson cycle.

The computational approach addressing the functional classification of FBA paralogous proteins is presented in this work for the first time. Indeed, the ProfileView method (Vicedomini, Bouly et al. 2019) has similarly been used for functional classification of other protein sequences across species. Here, we demonstrate that, based on it, one can classify paralogous sequences within a species by function, a problem which might be considered different from the classification problem formulated on homology. The results obtained on the FBA protein family open the door to new applications in protein sequence classification and their putative physiological role.

## Experimental procedures

### Cloning

Nuclear encoded amino acids sequence of *Chlamydomonas reinhardtii* Fructose-bisphosphate aldolase 3 (Cre05.g234550, UniRef100 entry Q42690) was analyzed by TargetP2.0 (Emanuelsson, Brunak et al. 2007) (Almagro Armenteros, Salvatore et al. 2019), ChloroP (Emanuelsson, Nielsen et al. 1999) and Predalgo (Tardif, Atteia et al. 2012) to predict the transit peptide. The subsequent mature sequence of chloroplastic protein coding for amino acid 28 to 377 has been amplified by PCR from the *Chlamydomonas reinhardtii* EST index (Asamizu and Nakamura 2004). PCR product was then inserted into pET3d-His6 by restriction and ligation at 5’-Nde1 and 3’-BamH1 sites to obtain a pET3d-His6-CrFBA3 expression plasmid. Plasmid sequences were validated by Sanger sequencing.

### Protein expression and purification

Transformation of *Escherichia coli* strain BL21(DE3) Rosetta-2 pLysS (Novagen Merck, Darmstadt Germany) were made with a pET3d-His6-CrFBA3 plasmid. This strain was grown in 2YT medium supplemented with ampicillin (100 μg/mL) at 37 °C until the culture reached exponential phase at an optical density at 600 nm of 0.8. The culture medium was supplemented with 0.2 mmol/L of IPTG and a temperature shift to 15 °C for 19 hours was made. Culture was next centrifugated at 4,000 *g* for 10 minutes at 4 °C. The cell pellet was resuspended in a buffer composed of 20 mM Tris-HCl (pH7.9), 100 mM NaCl, 5% glycerol and lyzed at a pressure of 2400 bar in a CF high-pressure homogenizer (Constant Systems Ltd., Daventry United Kingdom). The total extract of proteins was then centrifuged for 20 min at 20,000 *g* and the soluble fraction loaded on an affinity chromatography with 2 ml of NiNTA resin (Sigma-Aldrich Merck, Darmstad Germany). The resin was washed with 20 ml of 20 mmol/L Tris-HCl (pH7.9) supplemented with 100 mM NaCl (buffer A) complemented with respectively 10 mmol/L and 20 mmol/L of imidazole. CrFBA3 was then eluted in 2 consecutive steps with buffer A complemented with 250 mmol/L and 500 mmol/L of imidazole. Fractions were then analyzed by SDS-PAGE on a 12% acrylamide gel revealed by Coomassie blue staining. Fractions containing pure CrFBA3 were pooled, concentrated by ultrafiltration on 5,000 MWCO filter units (Millipore Merck, Darmstadt Germany) until a final volume of 10 ml prior to be injected on a size exclusion column HiLoad 26/600 Superdex 200 pg (Cityva life sciences) in buffer A. Fractions containing pure CrFBA were pooled, concentrated by ultrafiltration on 5,000 MWCO filter units (Millipore Merck, Darmstadt Germany) to a final concentration of 9 mg.mL^−1^ that was measured by NanoDrop 2000 spectrophotometer (Thermo Fisher Scientific, Waltham MA USA) with theoretical Mw = 39,390 g.mol^−1^ and ε_280_ = 50,037 mol^−1^.L.cm^−1^.

### Protein crystallization and structure determination

Purified CrFBA3 was tested for crystallization at two concentration of 9 mg/ml and 4.5 mg/ml on commercial sparse-screening conditions (Qiagen, Hilden Germany) based on the work of Jancarik and Kim (Jancarik, Scott et al. 1991) with a mixture of 50 nL of protein and 50 nL of precipitant solution equilibrated against 40 μL of reservoir solution at 20 °C. Several conditions showed small crystals in presence of ammonium sulfate. Condition A4 of Qiagen “the classics” kit (5% (v/v) isopropanol, 2 M ammonium sulfate) was selected to be optimized in a 2D gradient from 0% to 10% isopropanol and from 2 M to 2.5 M of ammonium sulfate with 2 μl drops of protein/precipitant solution ratio 1/1 equilibrated against 1 mL of reservoir solution. Protein concentration was brought to 2.6 mg/mL to avoid formation of clustered crystals and monocrystals were grown in a solution of 5 % isopropanol and 2 M ammonium sulfate. Crystals were flash frozen in liquid nitrogen for diffraction experiment at Proxima-2 beamline of SOLEIL synchrotron (Saint-Aubin, France). A 99.41 % complete dataset at 2.36 Å resolution was obtained from 3600 images and indexed in the C2 space group, integrated, scaled and converted with XDSME (Legrand 2017). Structure was phased by molecular replacement with PHENIX (Adams, Afonine et al. 2010) (Adams, Afonine et al. 2011) PHASER-MR (McCoy, Grosse-Kunstleve et al. 2007) using a search model obtained from the Phyre2 server (Kelley, Mezulis et al. 2015). Eight CrFBA monomers were found in the asymmetric unit. Model was then refined by iterative cycle of manual building in WinCOOT (Emsley and Cowtan 2004) (Emsley, Lohkamp et al. 2010) followed by refinement with PHENIX.REFINE (Afonine, Grosse-Kunstleve et al. 2012) until completion of a structure passing MOLPROBITY (Chen, Arendall et al. 2010) evaluation with 99.5% residues in Ramachandran restrains RMS(bond) = 0.008, RMS(angles) = 0.93 and final Rwork = 0.2002, Rfree = 0.2429 (Table 1). Structure representations were drawn with PYMOL 2.4 (Schrodinger, New York USA).

### Structural data

Reflections and coordinates files of the final model are registered in the protein data bank under accession code 7B2N.

### Computational analysis

#### Generation of functional trees

ProfileView (Vicedomini, Bouly et al. 2019) takes as input a set of homologous sequences and a protein domain, and returns a classification of the sequences in functional subgroups together with functional motifs characterising the subgroups. The first main idea of ProfileView is to extract conserved patterns from the space of available sequences through the construction of many probabilistic models for a protein family that should sample the diversity of the available homologous sequences and reflect shared structural and functional characteristics. These models are built as conservation profiles (pHMM [eddy1998profile]).

A pHMM can either generate new sequences, which are similar to the sequences used for its construction, or it can evaluate how likely it would be for a given query sequence to be generated by that model. To do so, it associates the query with an absolute likelihood score, which allows scores associated with different queries to be meaningfully compared.

ProfileView exploits this feature of pHMM for functionally describing an arbitrary set of query sequences. Indeed, the second main idea of ProfileView is to use its library of probabilistic models to embed input sequences into a multidimensional representation space, where each dimension is associated with a distinguished probabilistic model. Namely, each input sequence to be classified receives a score from each pHMM expressing how close the pHMM is to the sequence. All the scores are recorded as a vector entry for the sequence. Protein sequences are thus represented as points on this space, and distances can be calculated between each pair. These distances do not reflect mere sequence identities, but rather capture functional aspects that would have been overlooked by simpler, sequence-related metrics. Lastly, the distances are used to generate a tree that functionally relates all the query sequences. The internal nodes of the tree are, whenever possible, annotated by representative pHMM and by functional motifs. We expect this tree to discriminate paralogs taking part in Calvin-Benson cycle, such as *C. reinhardtii* FBA3, from paralogs involved in other chemical pathways (FBA1, FBA2, FBA4).

The *C. reinhardtii* FBA3 protein sequence (UniProt code Q42690) contains only one protein domain which encompasses the most conserved part of the protein. It is the large Pfam domain Glycolytic PF00274. In order to generate ProfileView functional tree for the 2601 homologous FBA sequences and including all *C. reinhardtii* FBA paralogous sequences, we devised the following pipeline:

1. Selection of the sequences for the construction of the probabilistic models To generate the ProfileView library of probabilistic models for the Glycolytic PF00274 domain, we downloaded the Pfam “full” alignment (Pfam version 34 - https://pfam.xfam.org/family/Glycolytic). We favored this set of sequences because of their relatively limited number of gaps (high sequence length homogeneity), hand curation and uniform sampling of homologous sequences on their phylogenetic tree. Of the 3886 sequences, 29% and 22% of them come from Proteobacteria and Metazoa respectively, and thus are not related to the Calvin-Benson cycle. Only 16% of sequences come from Viridiplantae. Nonetheless, it is important to select sequences covering most diversity as possible, in order to construct the most diversified model space for functional discrimination. No filter based on taxonomic classification was used in this step of the construction.
2. Clustering of the selected sequences Sequences having more than 20% gaps associated with amino acids of the query sequence were removed from the set, in order to filter out sequences with missing domains. The remaining 2356 sequences were clustered using MMseqs2 with parameters --min-seq-id 0.65 -c 0.8. The parameters were chosen as in (Ugarte, Vicedomini et al. 2018) and on its observation that optimal performances are obtained with 1000-1500 sequences. The resulting set contains 1244 sequences and spans the original diversity of taxa (supplementary figure 5).
3. Generation of the pHMM library We used the ProfileView-build routine to generate a library composed of 1244 pHMM models. These models are compiled from an equal amount of automatically-generated alignments, containing a variable number of sequences gathered in Uniprot (Uniclust30 08/2018)._
4. Generating the functional tree To determine the functional differences between FBA paralogs in photosynthetic species, we considered the complete Pfam “uniprot” dataset (15967 sequences) and selected sequences to be classified in the ProfileView tree based on taxonomy: we selected taxa with Uniprot classification Cryptophyceae, Glaucocystophyceae, Rhodophyta, and Viridiplantae. In order to have the maximum number of available sequences in the tree, no length-nor sequence identity-based filtering was applied. We identified 2601 sequences (of which 96% are Viridiplantae, 3% Cryptophyceae, 2% Rhodophyta, and only 2 Glaucocystophyceae). Note that the overlap between the 1244 sequences which are “seed” for the model construction and the 2601 classified sequences amounts to 252 sequences.

## Acknowledgements

We thank Mirko Zaffagnini for critical reading of the manuscript. This work was funded by ANR grants CALVINDESIGN (ANR-17-CE05-0001), CALVINTERACT (ANR-19-CE11-0009) and LABEX DYNAMO (ProjetIA-11-LABX-0011). We thank the *Institut de Biologie Physico-Chimique* (CNRS FR 550) for access to crystallization facility. We acknowledge SOLEIL for provision of synchrotron radiation facilities at beamlines Proxima-1, Proxima-2, and SWING.

