## Supplementary figures for "Crystal structure of chloroplast fructose-1,6-bisphosphate aldolase from the green alga *Chlamydomonas reinhardtii*"

### Slide 1
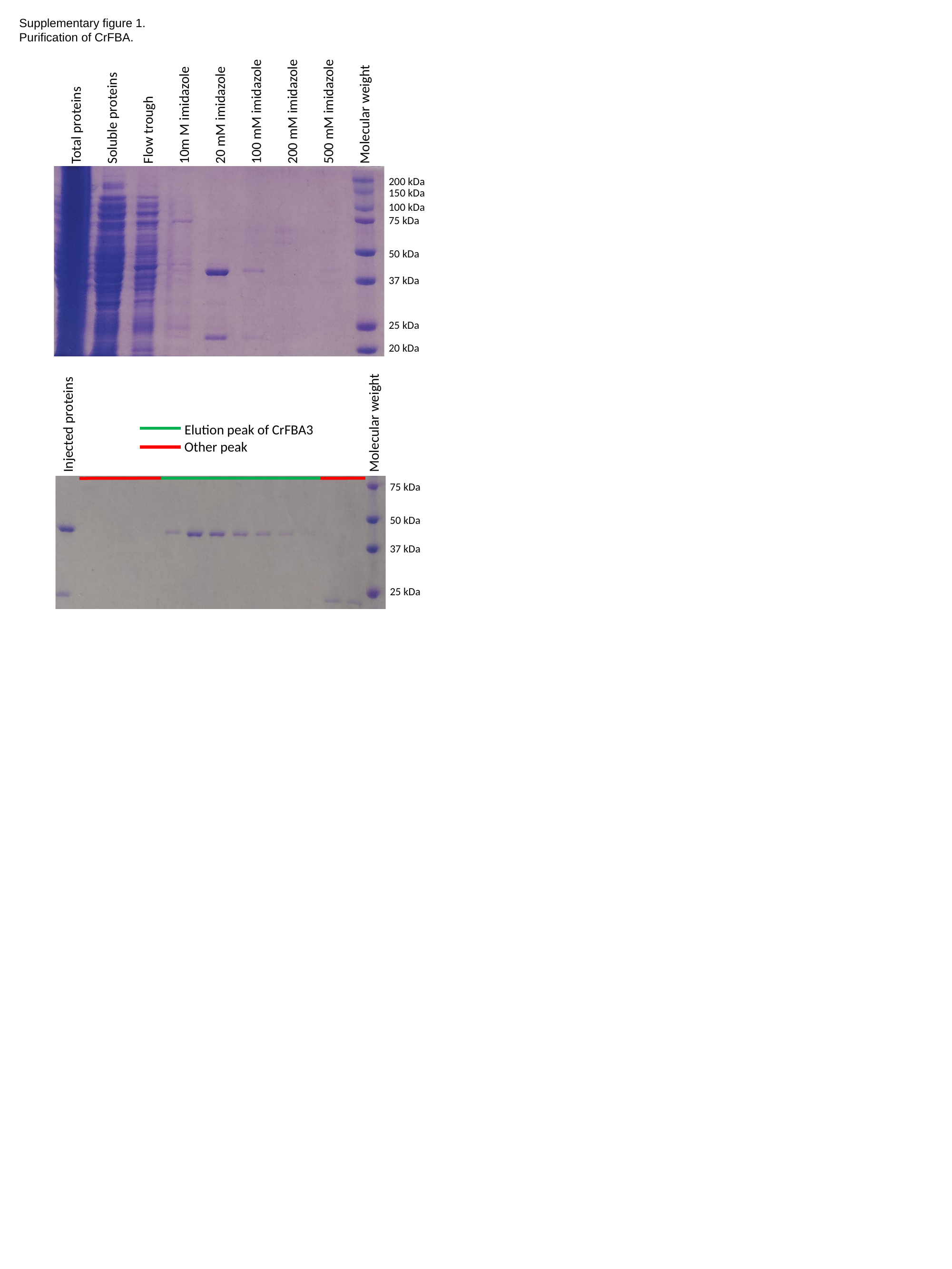

Supplementary figure 1.
Purification of CrFBA.
Total proteins
Soluble proteins
Flow trough
10m M imidazole
20 mM imidazole
100 mM imidazole
200 mM imidazole
500 mM imidazole
Molecular weight
200 kDa
150 kDa
100 kDa
75 kDa
50 kDa
37 kDa
25 kDa
20 kDa
Injected proteins
Molecular weight
Elution peak of CrFBA3
Other peak
75 kDa
50 kDa
37 kDa
25 kDa

### Slide 2
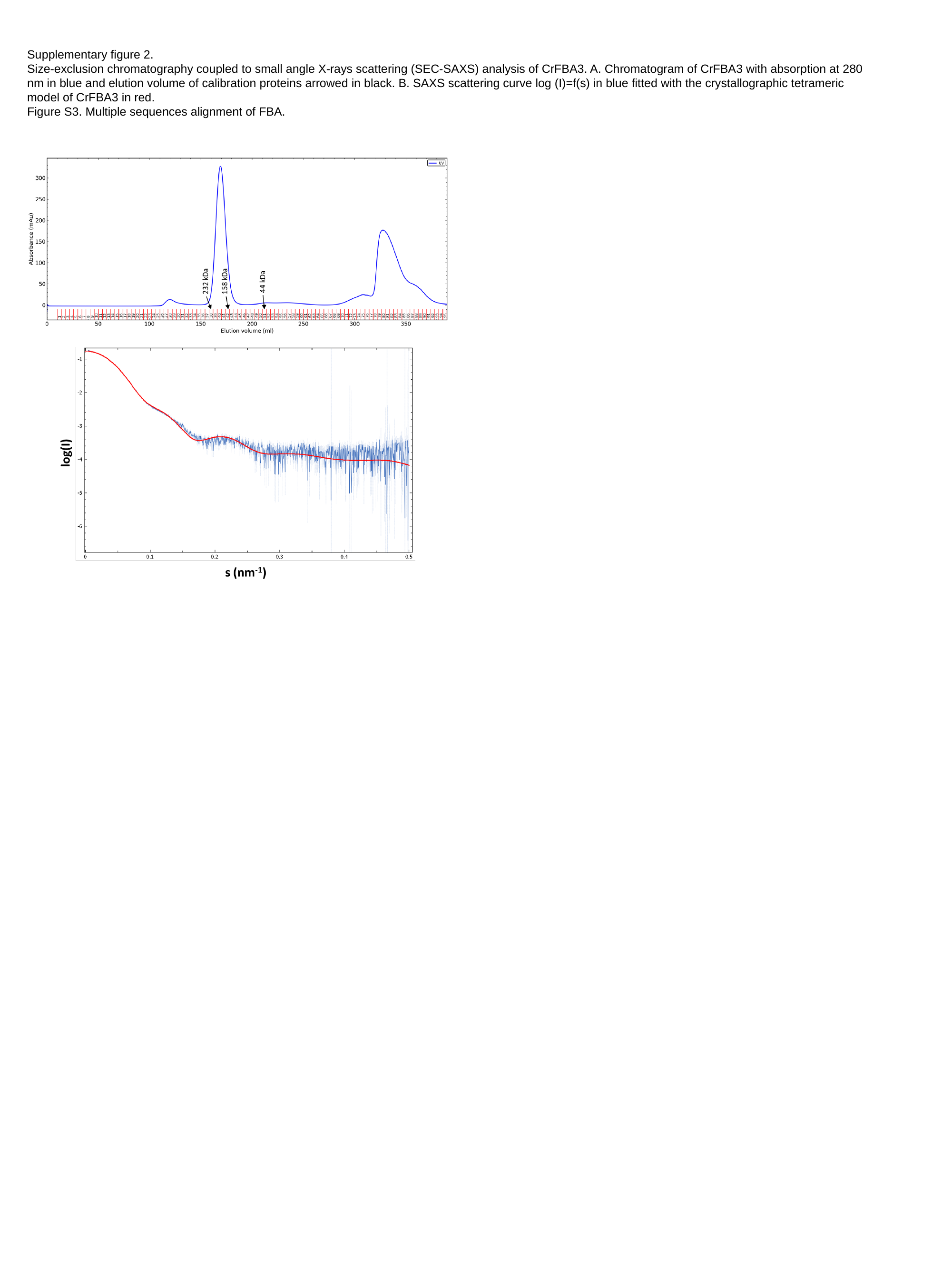

Supplementary figure 2.
Size-exclusion chromatography coupled to small angle X-rays scattering (SEC-SAXS) analysis of CrFBA3. A. Chromatogram of CrFBA3 with absorption at 280 nm in blue and elution volume of calibration proteins arrowed in black. B. SAXS scattering curve log (I)=f(s) in blue fitted with the crystallographic tetrameric model of CrFBA3 in red.
Figure S3. Multiple sequences alignment of FBA.

### Slide 3
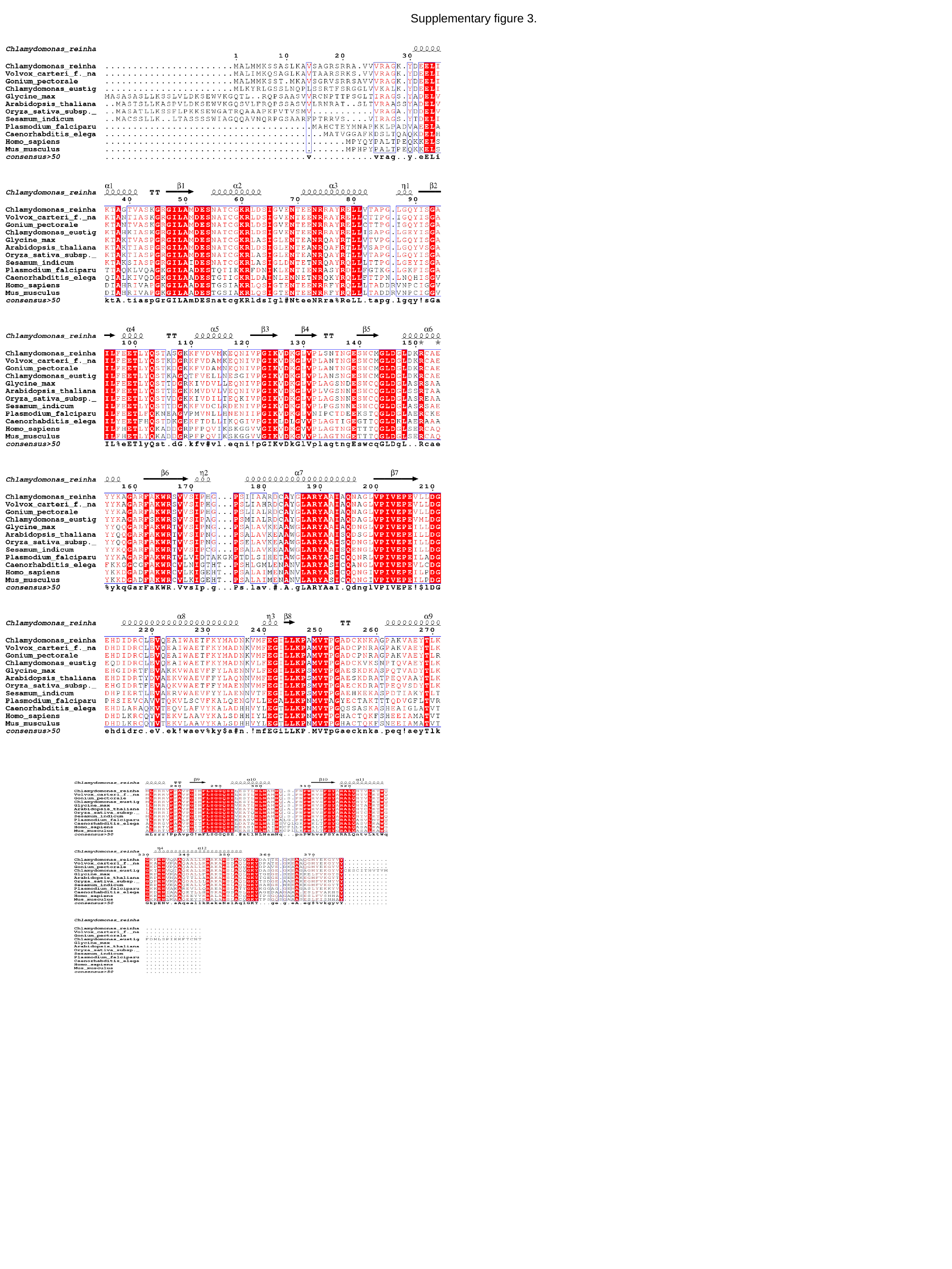

Supplementary figure 3.

### Slide 4
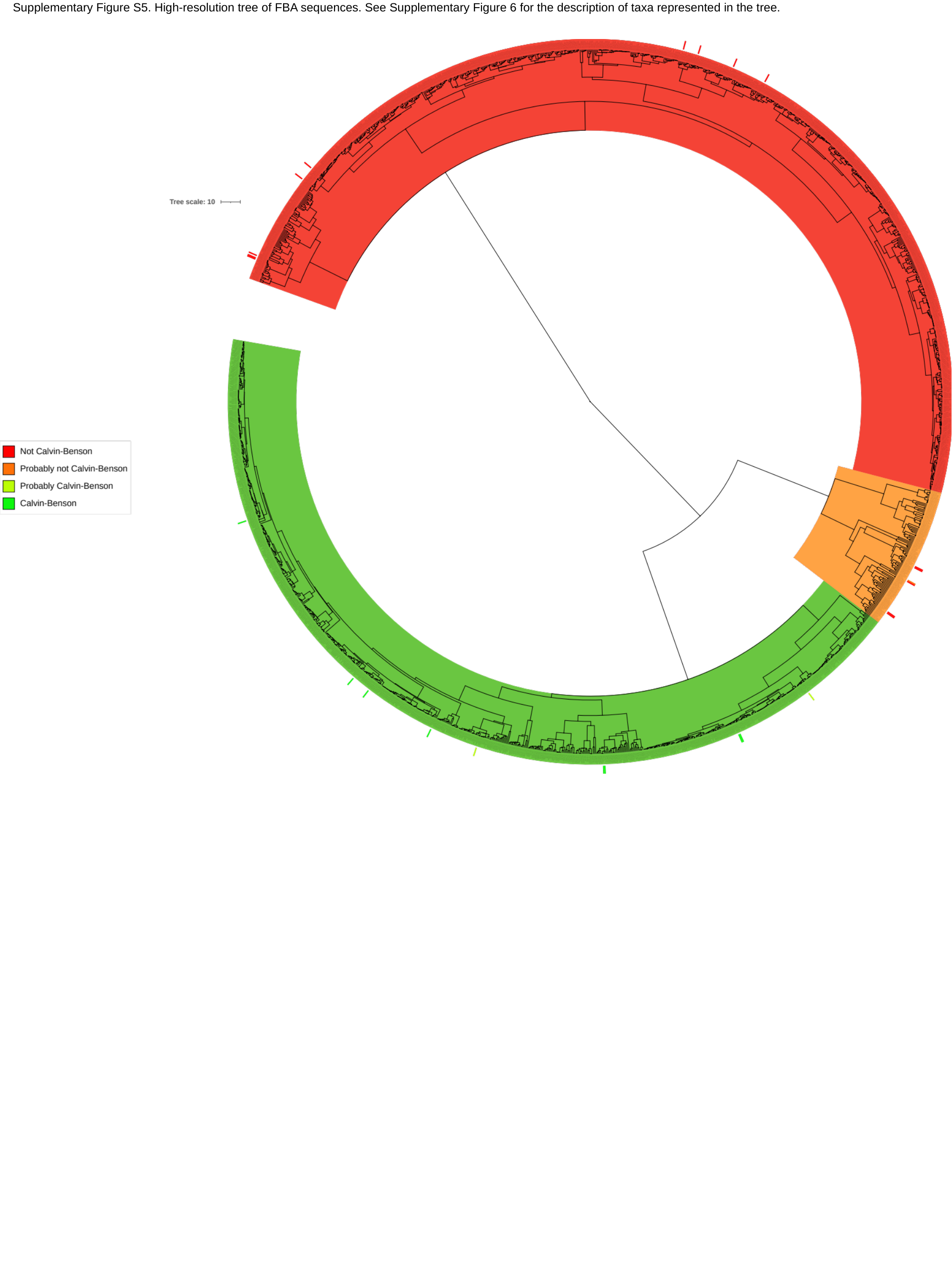

Supplementary Figure S5. High-resolution tree of FBA sequences. See Supplementary Figure 6 for the description of taxa represented in the tree.

### Slide 5
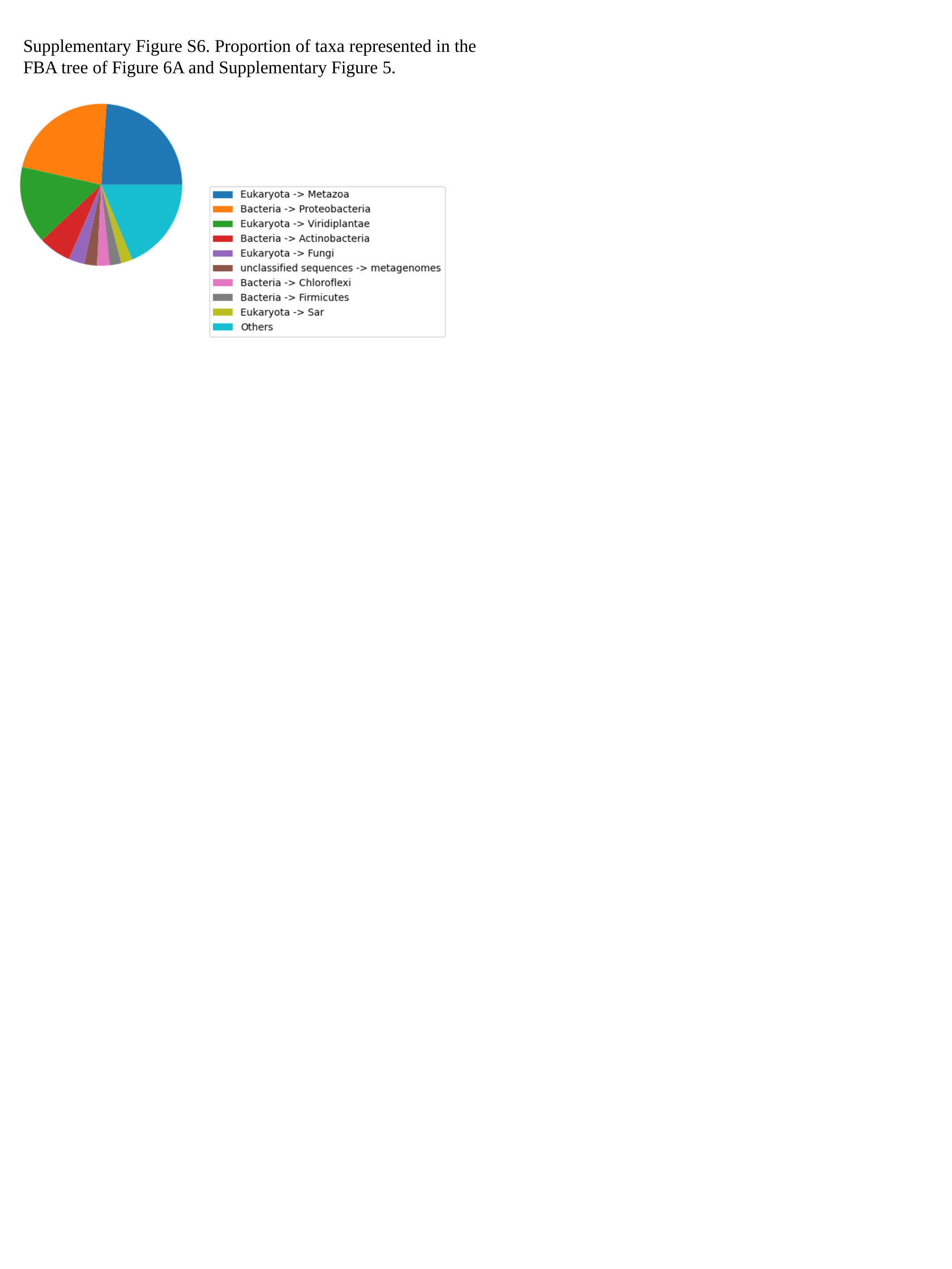

Supplementary Figure S6. Proportion of taxa represented in the FBA tree of Figure 6A and Supplementary Figure 5.
